# High Throughput Repurposing Screen Reveals Compounds with Activity Against *Toxoplasma gondii* Bradyzoites

**DOI:** 10.1101/2024.07.01.601569

**Authors:** Taher Uddin, Jing Xia, Yong Fu, Case W. McNamara, Arnab K. Chatterjee, L. David Sibley

**Affiliations:** Department of Molecular Microbiology, Washington University School of Medicine, St. Louis, MO, USA; Calibr at Scripps Research, La Jolla, California, USA

**Keywords:** High throughput screening, toxoplasmosis, chronic infection

## Abstract

*Toxoplasma gondii* causes widespread chronic infections that are not cured by current treatments due to inability to affect semi-dormant bradyzoite stages within tissue cysts. To identify compounds to eliminate chronic infection, we developed a HTS using a recently characterized strain of *T. gondii* that undergoes efficient conversion to bradyzoites in intro. Stage-specific expression of luciferase was used to selectively monitor growth inhibition of bradyzoites by the Library of Pharmacological Active Compounds, consisting of 1,280 drug-like compounds. We identified 44 compounds with >50% inhibitory effects against bradyzoites, including new highly potent compounds, several of which have precedent for antimicrobial activity. Subsequent characterization of the compound Sanguinarine sulfate revealed potent and rapid killing against in vitro produced bradyzoites and bradyzoites harvested from chronically infected mice. These findings provide a platform for expanded screening and identify promising compounds for further preclinical development against *T. gondii* bradyzoites responsible for chronic infection.

---

*Toxoplasma gondii* is a widespread parasite of animals that causes zoonotic infections in humans ^1^. Serological studies suggest that ∼ 1/3 of humans are chronically infected with *T. gondii* worldwide, although prevalence rates vary widely by geographic region ^2^. Human infections are caused by ingestion of undercooked meat, harboring tissue cysts, or ingestion of oocysts shed by infected cats ^3^. The acute stage of infection is predominated by fast growing tachyzoites that disseminate widely, including to sites of immune privilege like the brain, followed by differentiation into cysts harboring semi-dormant bradyzoites, which divide slowly and asynchronously ^4^. Although most infections are only mildly symptomatic and controlled by the immune system, they persist chronically and hence predispose individuals to subsequent reactivation if they become immunocompromised ^5^. The current standard of care treatment does not eradicate chronic infection, hence infected individuals remain at risk of reactivation for life.

Long known as a cause of congenital infection ^6^, recent studies highlight toxoplasmosis as a cause of ocular disease due to newly acquired infections in otherwise healthy adults ^7^. Frequent and severe outbreaks of ocular toxoplasmosis have been described in South America ^8^ and India ^9, 10^, and may occur in other localities. Notably, infection of healthy individuals in South America often leads to severe, recurrent ocular toxoplasmosis ^11^, with an estimated disease burden of 30 million individuals requiring treatment annually in Brazil alone ^12^. Additionally, prior toxoplasma infection has been linked to increased cognitive decline in Alzheimer’s Disease, although not all studies have conformed this association, likely due to the complexity of risk factors involved ^13^.

Current therapies for treatment of toxoplasmosis rely on inhibition of the folate pathway in the parasite ^14^. The standard of care therapy (i.e. sulfadiazine and pyrimethamine) inhibits the rapidly growing tachyzoite stage but has minimal activity on bradyzoites within tissue cysts and consequently does not eliminate chronic infection ^15^. Unfortunately, there are also significant adverse effects of this treatment regimen due to intolerance or allergic reactions ^16, 17^ and contraindication during the first two trimesters of pregnancy ^18^. Although drug resistance is not frequently encountered in treatment of toxoplasmosis, some isolates are naturally resistant to sulfonamides due to natural variants in dihydropteroate synthetase, or other molecular mechanisms, thus complicating treatment in some cases ^17^. Clindamycin and several macrolide antibiotics have also been shown to inhibit growth in vitro and in animal models ^19, 20^ and such antibiotics have been used to treat toxoplasmosis in humans ^21^. However, these compounds are not specific to the parasite and disrupt the normal endogenous microbiota leading to possible emergence of pathobionts like *Clostridium difficile* ^22^.

A review of the literature of compounds that are clinically approved in humans or animals identified several compounds that inhibit parasite growth in vitro and/or in murine models of toxoplasmosis ^21^. For example, repurposing of guanabenz, an FDA-approved drug that interferes with translation, showed activity against acute and chronic toxoplasmosis in mice ^23^, although this affect was dependent on the strain of mouse ^24^, and treatment did not eliminate cysts, resulting in rebound after discontinuation ^25^. A repurposing screen for inhibition of tachyzoite growth in vitro using the Tocriscreen Total Library, which consists of 1,280 biologically active small molecules with confirmed molecular targets, identified multiple compounds that affect dopaminergic and estrogen signaling, including tamoxifen that was shown to act by upregulating xenophagy to restrict parasite growth and cause clearance ^26^. Additionally, a number of new investigational compounds have shown an ability to block tachyzoite growth in vitro and during acute infection in animal models, including several that have activity against bradyzoite growth or chronic infection ^17^.

The majority of efforts to identify new compounds with activity against *T. gondii* have focused on in vitro assays using tachyzoite growth as a readout ^17, 21^. Differentiation of bradyzoites in vitro can be achieved by treatment with stress, such as high pH ^27^; however, the stages that develop under stress often continue to express tachyzoite traits ^28, 29^,thus complicating screening efforts. Nonetheless, combining dual promoters to drive firefly luciferase in the cytosol of bradyzoites and Nanoluc luciferase (nLuc) that was engineered to be secreted into the cyst matrix, allowed evaluation of compounds for selective activity against chronic stages ^30^. Another promising development is the recent development of an in vitro system for development of bradyzoites using a specialize KD3 muscle cell line where spontaneous differentiation occurs ^31^. Consistent with the lack of available treatments on tissue cysts in chronically infected mice, treatment with pyrimethamine and/or sulfadiazine was not effective in restricting the growth of mature bradyzoites formed in vitro in this system ^31^. Hence, this system could provide a useful platform for testing compounds for activity against bradyzoites. However, this system requires the use of a specialized culture system and the treatment and recovery phases needed for evaluation takes ∼ 50 days, complicating its use for screening.

As an alternative, we recently described a type II strain called Tg68 that has a high propensity to differentiate into bradyzoite in vitro under condition of stress that include high pH or cultivation in high glutamine, low glucose, that forces metabolism based on glutaminolysis ^32^. Unlike other type II strains that undergo partial differentiation, Tg68 forms fully mature bradyzoites without associated breakthrough of tachyzoites following stress induction in vitro ^32^. Here, we engineered this strain to express Firefly luciferase (Fluc) under the control of a constitutive promoter, and separately generated a line expressing nLuc under the control of a bradyzoite promoter. We used these reporter lines to develop a high throughput screen (HTS) and used it to evaluate the Library of Pharmacological Active Compounds (LOPAC). Several compounds with activity against both stages were identified, providing proof of concept for further HTS projects designed to find new treatments for chronic toxoplasmosis.

## Results and Discussion

To facilitate HTS using the Tg68 strain, we generated a clonal line of Tg68 constitutively expressing firefly-luciferase (Fluc) under the p*TUB1* promoter (**Figure 1A**). The pTub1:Fluc plasmid also contained a resistant DHFR cassette that was integrated into the genome after electroporation followed by selection with pyrimethamine. Following passage in HFF cells, a cloned line constitutively expressing Fluc was isolated and is referred to as Tg68-pTub1:Fluc. Confluent HFF cells grown in 384-well plates were infected with Tg68-pTub1:Fluc tachyzoites at 2×10^4^ parasites per well, and the infection was allowed to proceed for 72 hr under 5% CO_2_ at 37°C (**Figure 1B**). Comparison of Fluc activity between day 3 and day 0 (4 hr postinfection) increased significantly, representing growth of the parasite (**Figure 1C**). In addition, we analyzed Fluc expression from replicate 14 wells, across 5 plates and determined the Coefficient of Variation (CV) for positive wells was 7±3% (**Table S1**). Addition of BRD7929, which targets parasite phenylalanine tRNA synthetase ^33^, completely inhibited parasite growth at a concentration of 10 μM (**Figure 1D**). Comparison of the luciferase signal from untreated and treated wells from 5 plates demonstrated an average Z’ value of 0.77±0.11 (**Table S1**). Additionally, the Tg68-pTub1:Fluc line demonstrated comparable sensitivity to BRD7929 and atovaquone compared to the reference type II strain TgMe49-Fluc (**Figure 1D**). Taken together, these findings indicate that Tg68-pTub1:Fluc provides a robust readout for HTS of compounds against tachyzoite growth.

**Figure 1.**
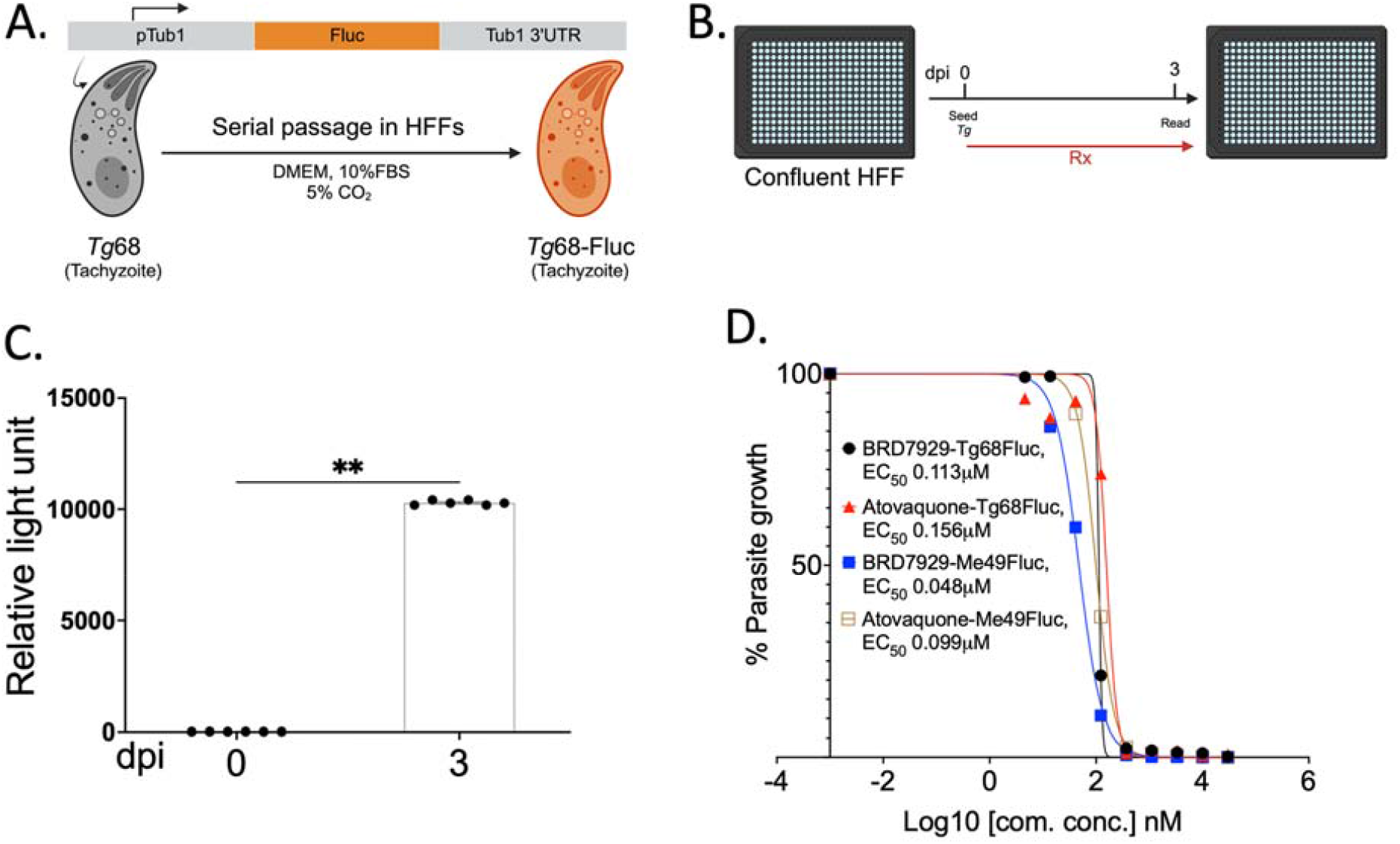
Development of a High Throughput Screening assay for growth inhibition of Tg68 tachyzoites. (A) A tachyzoite-specific firefly-luciferase (Fluc) reporter strain of Tg68 was generated using the pTUB1 promoter. (B) Confluent HFF cells in 384-well plates were infected with Tg68 tachyzoites and luciferase activity was performed on day 3. (C) Replicate of 12 wells from six independent plates was analyzed for luciferase activity. Fluc expression increased more than 1,000-fold and was found significantly higher at day 3 hr (72hr) compared to day 0 (4 hr postinfection). Data accumulated from 12 wells, across 6 plates. Mann-Whitney test, *P* < 0.02. (D) EC_50_ determination for Tg68-pTub1:Fluc and ME49-Fluc treated with serial dilutions of BRD7929 or Atovaquone. All EC_50_ values are presented as the mean of three biological replicates (n = 3)

To expand the use of Tg68 for HTS of compounds with activity against bradyzoites, we generated a clonal line expressing nLuc under control of the p*BAG1* promoter, which is strongly upregulated upon differentiation ^34^. The pBAG1:nLuc plasmid also contained a resistant DHFR cassette that was integrated into the genome after electroporation followed by selection with pyrimethamine. Following serial passage in HFF cells, a positive clone was isolated and referred to as Tg68-pBAG1:nLuc. (**Figure 2A**). In previous studies, we have shown that the Tg68 strain undergoes highly efficient differentiation to bradyzoites in alkaline media at low CO_2_ in vitro and that it retains a mature bradyzoite profile even at relatively high multiplicities of infection ^32^. Tg68 also undergoes efficient differentiation when grown in glucose free medium supplemented with 10 mM glutamine, a process that forces glutaminolysis for energy production via the mitochondrion ^32^. Confluent HFF cells in 384-well plates were infected with pBAG1:nLuc parasites ( 3x10^3^ parasites per well) for 2 hr, followed by washing and shifting to alkaline or glutamine media. Cultures were maintained for 10 days under CO_2_-free conditions (ambient air), with media changes on day 3 and 6. On day 6, compounds were added and on day 10 luciferase assays were performed (**Figure 2B**). Expression of nLuc was very low in cultures of tachyzoites and in the initial culture conditions at day 0 (4 hr postinfection) in normal or differentiation media (**Figure 2C**). Expression of nLuc increased dramatically by over 4 logs by day 3 and continued to increase significantly at day 6 and at day 10 (**Figure 2C**). On day 10, analysis of luciferase values from 14 wells showed an optimal Coefficient of Variation (CV) of 10±2% for alkaline media and 7±2% for glutamine media, indicating suitability for high-throughput screening (HTS) (**Table S1**). Treatment with BRD7929 at a concentration of 10 μM completely inhibited growth of the parasites and comparison of nLuc activity from treated and untreated wells revealed a Z’ value of 0.67±0.05 for alkaline media and 0.76±0.06 for glutamine media (**Table S1**). We also compared the potency of BRD7929 and atovaquone under both conditions that induced bradyzoite differentiation (**Figure 2D**). Both compounds showed a reduction in EC_50_ values under glutamine differentiation when compared to alkaline conditions, although this was much more dramatic for atovaquone that showed a > 30-fold shift (**Figure 2D**). The greater potency of compounds in glutamine medium may reflect a decreased ability of the parasite to generate energy stores from glutaminolysis vs. glycolysis as suggested previously by the modest growth defects in knockouts of *T. gondii* hexokinase ^35^ and glucose transporter 1 ^36^. Consequently, the use of glutamine medium for bradyzoite development may preferentially reveal compounds that act on the mitochondrion, as is the case for atovaquone that inhibits the bc1 complex ^37^.

**Figure 2.**
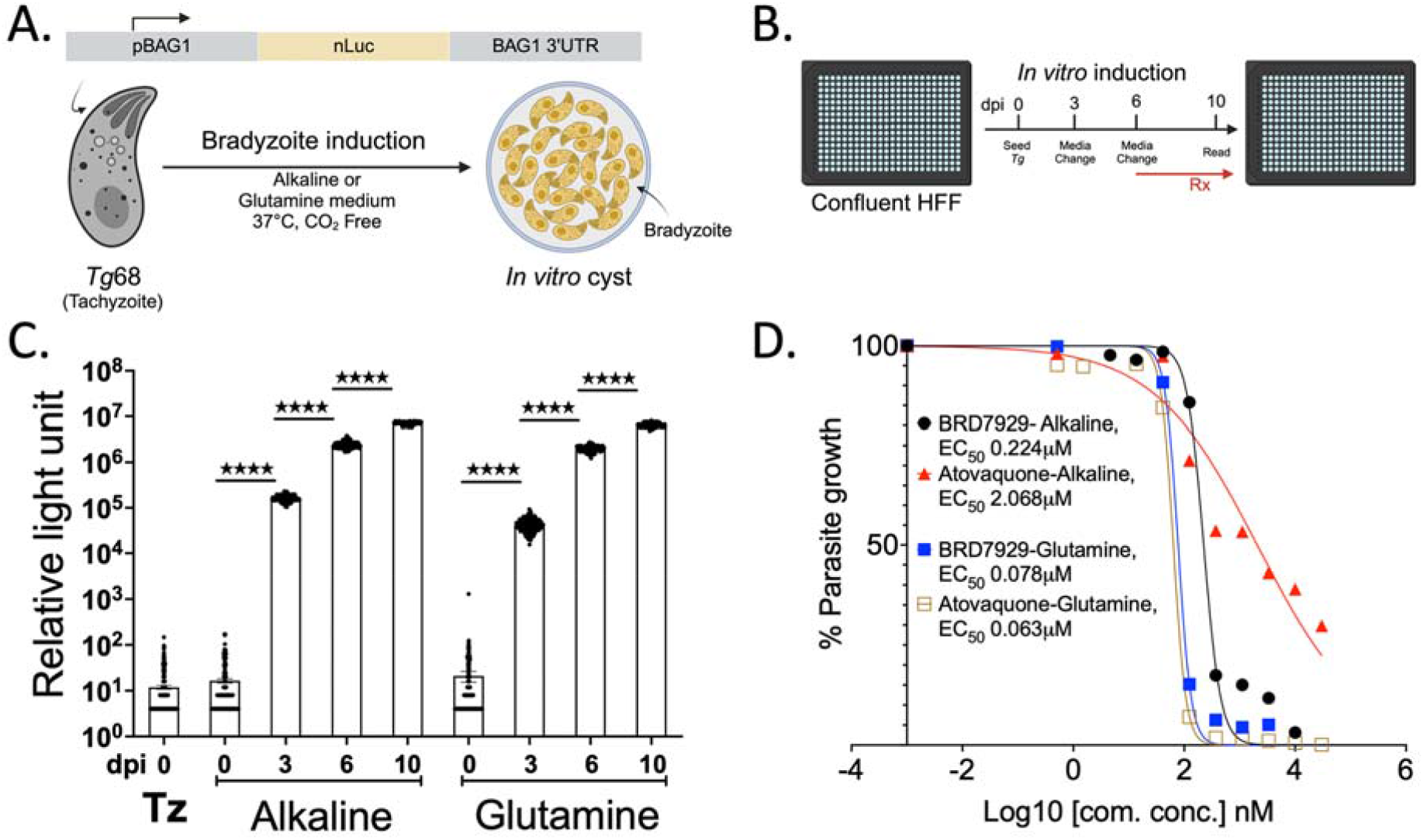
Development of a High Throughput Screening assay for growth inhibition of Tg68 bradyzoites. (A) A bradyzoite-specific Nanoluc luciferase (nLuc) reporter strain of Tg68 was generated using the p*BAG1* promoter. Parasites were grown in alkaline medium (pH 8.2), CO_2_ free or in the absence of glucose supplemented with glutamine, which stimulates in vitro development of bradyzoites. (B) Confluent HFFs in 384-well plates were infected with Tg68 tachyzoites for two hr, washed and then cultured either in D10 under normal conditions (Tz) or switch to alkaline or glutamine conditions to induce bradyzoites. The cultures were then maintained under CO_2_-free conditions for 10 days with media changes on day 3 and 6, with a compound treatment at day 6 and readout at day 10. (C) Luciferase signals from tachyzoites (Tz) harvested at day 0 (4 hrs postinfection) postinfection or bradyzoites induced for different times (day 0 (4 hr) to day 10) by culture in alkaline or glutamine media. Comparisons between sequential time points using Mann Whitney test, **** P < 0.0001(D) Determination of EC_50_ values for BRD7929 and atovaquone treatment of in vitro induced bradyzoites culture in alkaline or glutamine media. All EC_50_ values are presented as the mean of four biological replicates (n = 4).

The LOPAC library (1157 of 1280 total compounds) was plated in 384 well format and tested for parasite growth inhibition in duplicate using a single concentration of 10 μM in the tachyzoite and both alkaline and glutamine-induced bradyzoite assays. Growth inhibition was averaged from the two replicates and plotted as a Venn diagram summarizing the outcome of each of the three assays (**Figure 3A**). A total of 27 compounds showed selective inhibition of tachyzoite growth without affecting growth in the other assays (**Figure 3A**). The largest number of compounds was identified in the glutamine induced bradyzoite assay, perhaps reflecting the metabolic liability of this growth condition (**Figure 3A**). In total, 21 compounds that inhibited parasite growth in all three assays by 50% or more were identified as Primary Hits (**Figure 3A**). Furthermore, we identified 9 compounds that inhibited bradyzoite growth in both assays by 50% or more but not tachyzoite growth (**Figure 3A**). Finally, we identified 14 compounds that specifically inhibited glutamine-induced bradyzoite growth by ≥ 80% but were not effective in growth inhibition in the other two assays (**Figure 3A**). Robust validation of the within plate and between replicates was performed through collective calculation of the coefficient of variation (CV) and Z’ analysis of assay plates (**Table S2**). Of the 44 Primary Hit compounds defined above (**Figure 3B**), 36 compounds were available in quantity at Calibr and were chosen for further testing (**Table S3**). These 36 compounds were tested in a dilution series and 9 compounds were found to have EC_50_ values of ≤ 2 µM in either tachyzoite and/or bradyzoite assays, thus defining a set of Top Hits (**Table S3, Figure 3B**).

**Figure 3.**
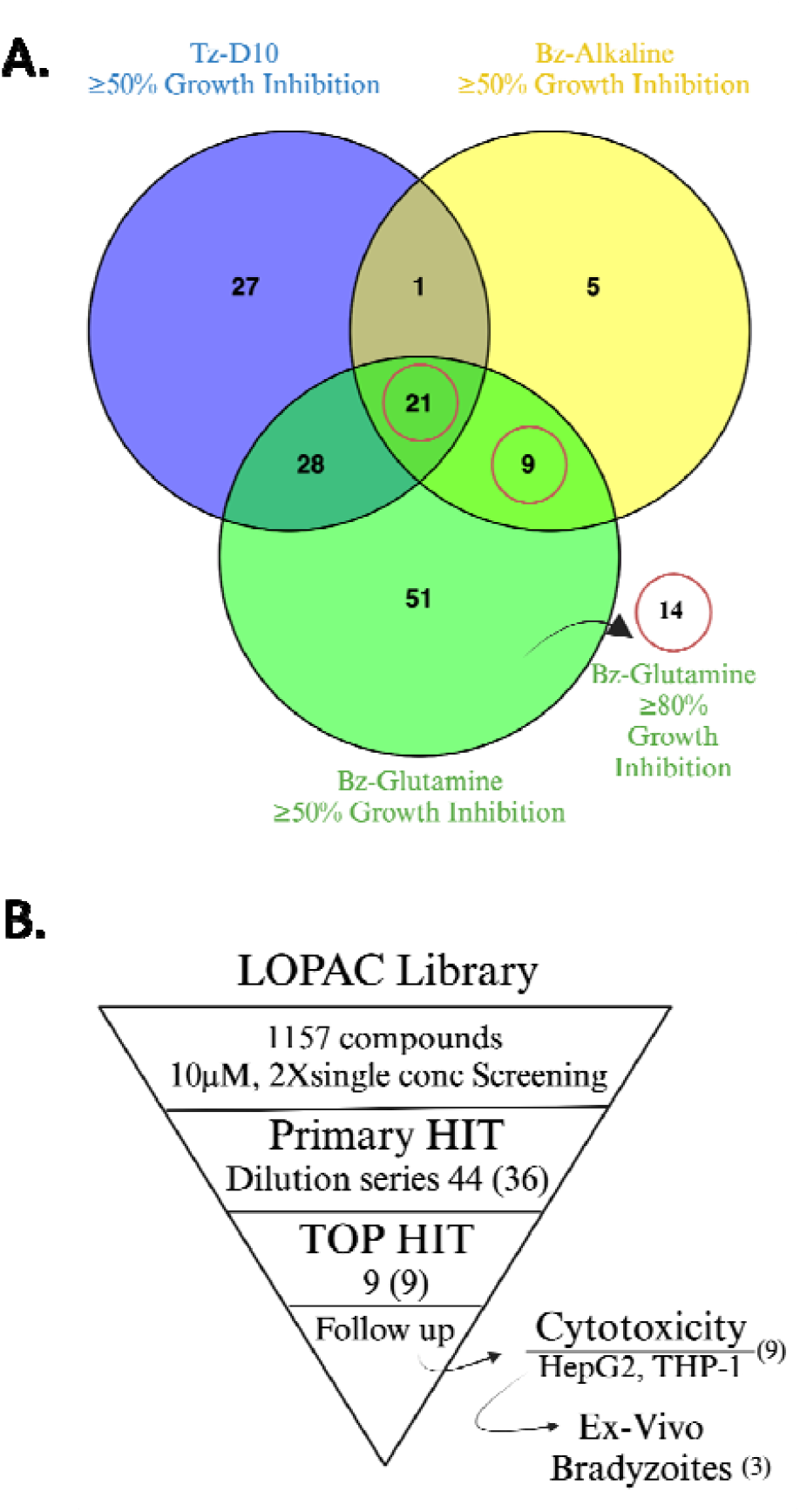
Summary of LOPAC screening for growth inhibition of *T. gondii*. (A) Venn diagram showing the number of compounds with ≥ 50% of growth inhibition at 10 μM in each of three growth assays. Tz = tachyzoite growth assay, Alk = alkaline induced bradyzoite growth assay, Gln = glutamine induced bradyzoite growth assay. Red circled numbers indicate the selection criteria for Primary Hits. (B). Summary of LOPAC screen and prioritization of Hits for follow up. Of 44 Primary Hits, 36 were available for dilution series to determine EC_50_ values. Top 9 Hits were available for further biological testing.

We were able to source the 9 Top HIT compounds from commercial sources. The 9 available compounds were tested for cytotoxicity against HepG2 and THP-1 cell lines, along with a control drug, atovaquone (**Figure 3B**). Most compounds exhibited a favorable selectivity index (SI), which is a measure of the compound’s EC_50_ against the parasite compared to the CC_50_ against host cells (**Table 1**). However, a few compounds, namely Idarubicin hydrochloride and MS012, showed high toxicity, while Auranofin and JFD00244 showed modest toxicity towards the cell lines. Auranofin is a gold containing compound that has been approved for treatment of rheumatoid arthritis and it has previously been reported to have activity against several parasites ^38^. Consistent with this profile, auranofin has previously been shown inhibit replication of the type I RH strain in vitro, reduce infection in a chicken embryo model ^39^and reduce the burden of cysts in chronically infected mice ^40^. However, the high level of growth inhibition for HepG2 and THP-1 cells treated with auranofin in the present study would appear to limit the potential of this compound for further clinical development. Additionally, Brefeldin A, which blocks ER to Golgi transport showed a favorable SI in HepG2 cells but not in THP-1 cells (**Table 1**), suggesting that block of protein export has very different consequences on host cell growth in different lineages. As a result, these compounds were excluded from further assays due to their undesirable cytotoxic effects. Emetine dihydrochloride hydrate

**Table 1.**
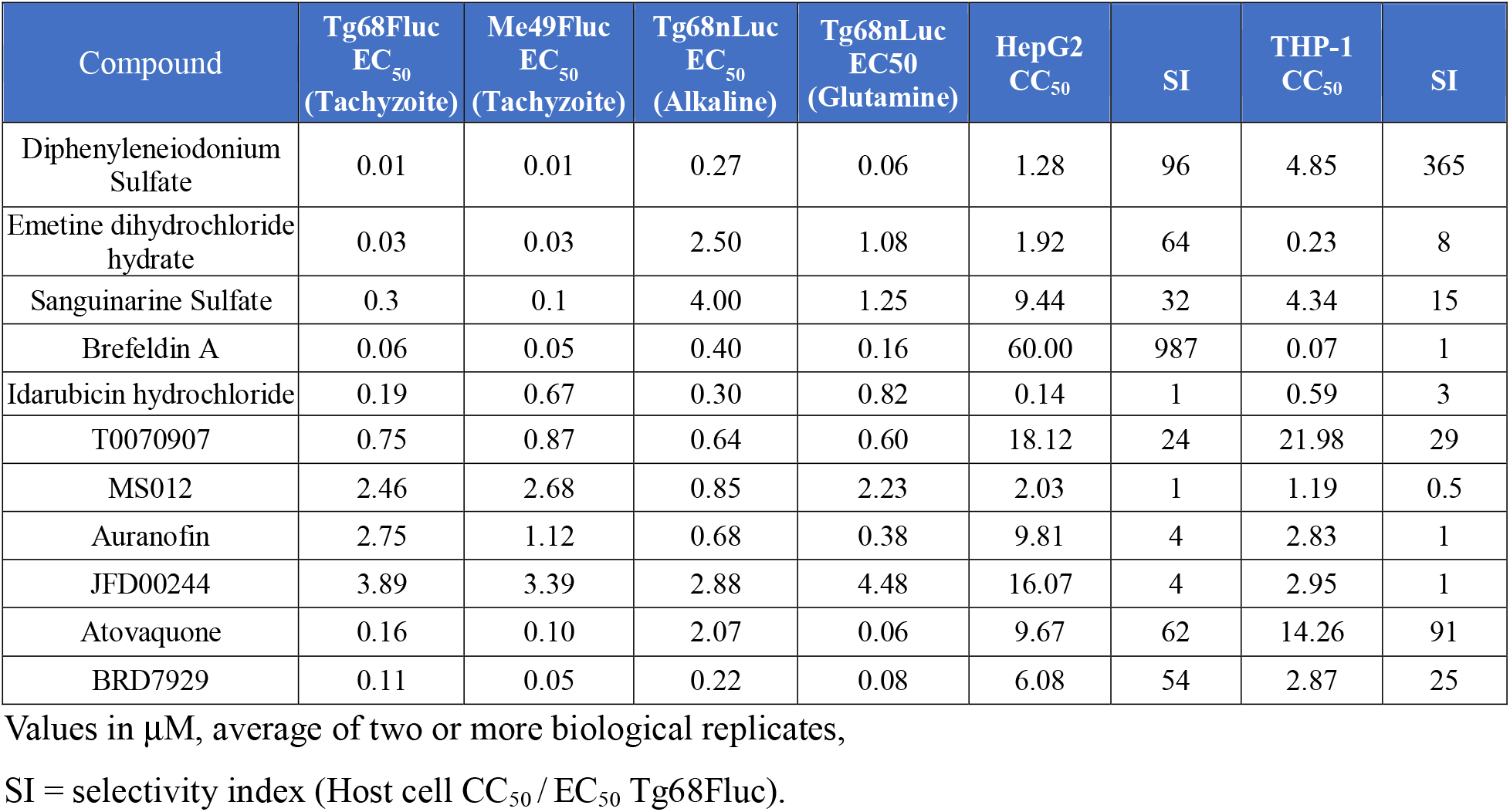
Potency and selectivity of Top Hits

| Compound | Tg68Fluc<br>EC<sub>50</sub><br>(Tachyzoite) | Me49Fluc<br>EC<sub>50</sub><br>(Tachyzoite) | Tg68nLuc<br>EC<sub>50</sub><br>(Alkaline) | Tg68nLuc<br>EC<sub>50</sub><br>(Glutamine) | HepG2<br>CC<sub>50</sub> | SI | THP-1<br>CC<sub>50</sub> | SI |
| --- | --- | --- | --- | --- | --- | --- | --- | --- |
| Diphenyleneiodonium Sulfate | 0.01 | 0.01 | 0.27 | 0.06 | 1.28 | 96 | 4.85 | 365 |
| Emetine dihydrochloride hydrate | 0.03 | 0.03 | 2.50 | 1.08 | 1.92 | 64 | 0.23 | 8 |
| Sanguinarine Sulfate | 0.3 | 0.1 | 4.00 | 1.25 | 9.44 | 32 | 4.34 | 15 |
| Brefeldin A | 0.06 | 0.05 | 0.40 | 0.16 | 60.00 | 987 | 0.07 | 1 |
| Idarubicin hydrochloride | 0.19 | 0.67 | 0.30 | 0.82 | 0.14 | 1 | 0.59 | 3 |
| T0070907 | 0.75 | 0.87 | 0.64 | 0.60 | 18.12 | 24 | 21.98 | 29 |
| MS012 | 2.46 | 2.68 | 0.85 | 2.23 | 2.03 | 1 | 1.19 | 0.5 |
| Auranofin | 2.75 | 1.12 | 0.68 | 0.38 | 9.81 | 4 | 2.83 | 1 |
| JFD00244 | 3.89 | 3.39 | 2.88 | 4.48 | 16.07 | 4 | 2.95 | 1 |
| Atovaquone | 0.16 | 0.10 | 2.07 | 0.06 | 9.67 | 62 | 14.26 | 91 |
| BRD7929 | 0.11 | 0.05 | 0.22 | 0.08 | 6.08 | 54 | 2.87 | 25 |

Values in $\mu\text{M}$ , average of two or more biological replicates,
SI = selectivity index (Host cell CC<sub>50</sub> / EC<sub>50</sub> Tg68Fluc).

Although Tg68 undergoes efficient conversion to bradyzoites in vitro, with a transcriptional profile that resemble in vivo bradyzoites ^32^, it may still lack some of the features of mature tissue cysts. Hence, we tested select Top Hits, along with several reference compounds, against tissue cysts that were harvested from chronically infected mice (**Figure 4A**). For these assays, we used the ME49 EW strain, which produces high numbers of cysts in vivo ^41^. As shown in **Table 1**, the ME49 strain has a very similar sensitivity to the Top Hits and reference compounds, thus validating the choice of strain. Compounds were tested using continuous exposure of ex vivo bradyzoites to 3XEC_90_ as measured on tachyzoites. Alternatively, ex vivo bradyzoites were treated at 3XEC_90_ for only 4 hr followed by washout to ascertain how irreversibly they might act (**Figure 4A**). All three of the reference compounds BRD7929, atovaquone, and pyrimethamine showed potent inhibition of bradyzoites when used continuously (**Figure 4B**). Although atovaquone and BRD7929 are active on bradyzoites (**Table 1**), the activity of pyrimethamine in this continuous treatment assay is likely because it inhibits the outgrowth of tachyzoites. Consistent with this specificity, pyrimethamine was largely ineffective when used for only 4 hr, indicating it has minimal effects on bradyzoites present at the start of the assay (**Figure 4B**). Similar to pyrimethamine, testing of diphenyleneiodonium and T0070907 in the ex vivo bradyzoite assay revealed that they only work when used in continuous treatment, suggesting their activity is static rather than cidal (**Figure S1**). Diphenyleneiodonium is an inhibitor of NADPH oxidase that separately induces oxidative stress ^42^. Previous studies have shown diphenyleneiodonium inhibits growth of *T. gondii* tachyzoites in ARPE-19 cells through the production of ROS ^43^. In a separate study, it also showed activity against *P. falciparum* with an EC_50_ of 0.06 nM ^44^, and it has also been shown to have broad spectrum antibacterial activity ^45^. T0070907 is an inhibitor of peroxisome proliferator activator receptor γ that induces G2/M arrest and thus has activity against cancer cells ^46^. This nuclear hormone pathway is not conserved in *T. gondii*, and this compound has not been described to have anti-microbial activity previously, so the potential mechanism of action is uncertain.

**Figure 4.**
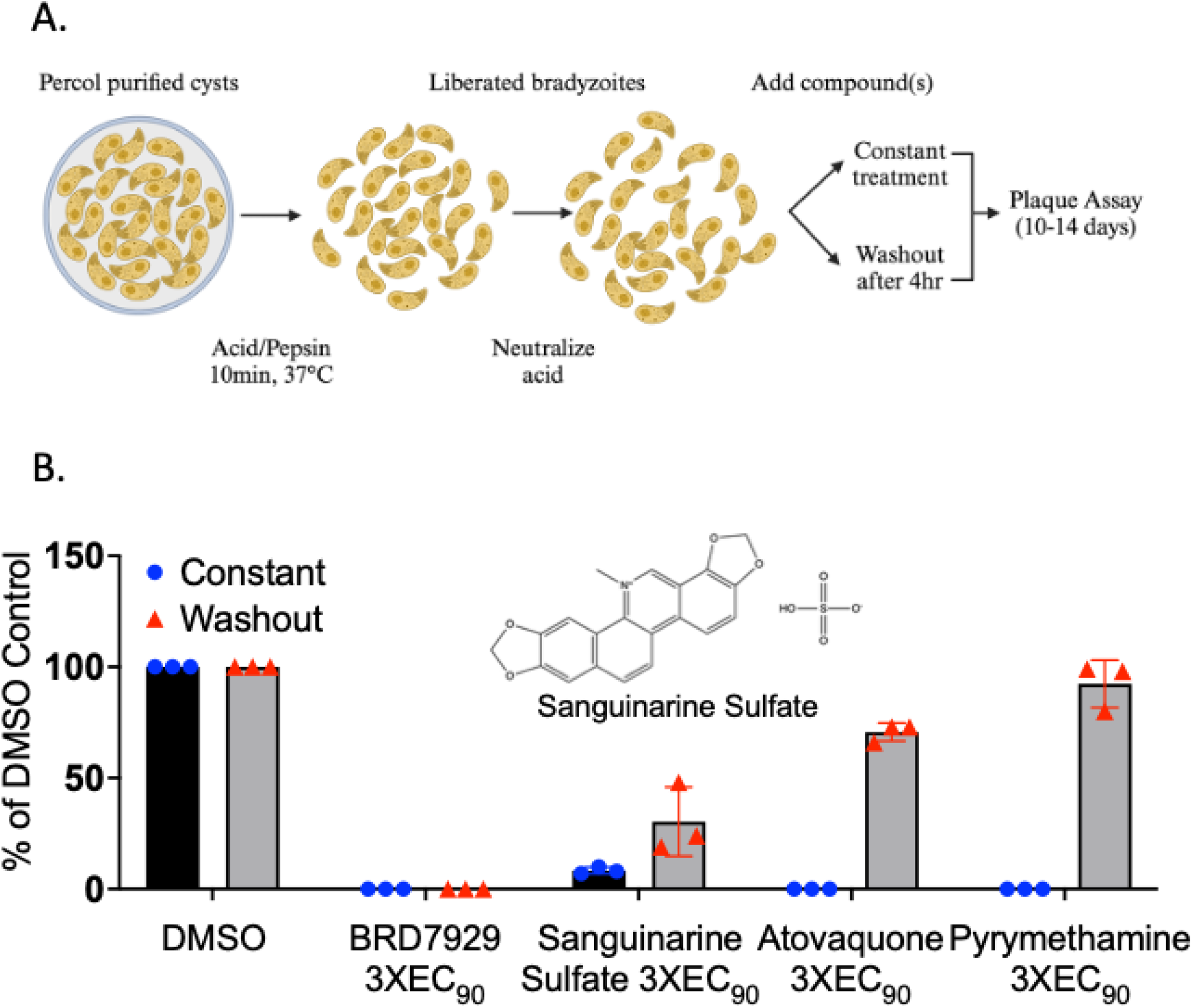
Testing of compounds for activity against ex vivo bradyzoites. (A) Schematic for isolation of bradyzoites from chronically infect mice and testing in vitro during continuous treatment or after 4 hr treatment and washout. (B) Testing of the Top Hit Sanguinarine and reference compounds in continuous treatment vs. 4 hr treatment and washout. Values are determined from plaque counts after 10-14 days of outgrowth and are normalized to the DMSO control for each condition. Compounds were used at 3XEC_90_ based on sensitivity of Tg68Fluc tachyzoites (**Table S3)**. Data from 3 biological replicates, bar graph represent the average of percentage of number of plaques in the treatment group normalized to DMSO control.

Atovaquone showed partial inhibition in the washout ex vivo bradyzoite assay, and consistent with previous studies showing it is partially active in reducing cysts numbers during chronic infection in vivo ^47^. BRD7929 was highly effective in preventing the outgrowth of bradyzoites even when removed after 4 hr (**Figure 4B**), consistent with previous findings ^33^. Testing of the Top Hit Sanguinarine sulfate revealed potent activity in the ex vivo bradyzoite wash out assay, thus confirming it has activity against bradyzoites as well as tachyzoites (**Figure 4B**). Sanguinarine sulfate is a natural product produced by the opium poppy and several other plants and it consists of a benzoquinoline alkaloid ^48^. Sanguinarine has anti-inflammatory, anti-tumor and antimicrobial activities and it is thought to act on numerous signaling pathways in human cells ^49^. Sanguinarine has been described to be toxic to host cell, thought to be due to its action on Na^+^/K^+^ ATPase, although other mechanisms have also been described ^48^. Although we did not measure appreciable cell toxicity in our assays, other studies have indicated sanguinarine induces apoptosis or blocks cell growth ^49^. Additionally, the LD_50_ in mouse is 18 mg/kg by i.p. injection ^49^, likely limiting its use in efficacy testing for toxoplasmosis. Emetine is a natural product that binds to ribosomes and blocks translation, and studies of the structure of *Giardia* ribosomes have recently refined the binding mode and mechanism of action ^50^. Emetine has also been shown to be potent against *Plasmodium falciparum* in vitro ^51^and against *Trypanosoma brucei* and *Trypanosoma cruzi*, although it displayed poor selectivity relative to host cells ^52^. Although we did not observe toxicity in short term CC_50_ assays, it was toxic to monolayers in longer term culture, preventing us from testing its effects against ex vivo bradyzoites. Advancing emetine as a treatment for parasitic infections would also be compromised by its cardiotoxicity ^53^.

## Conclusions

The newly discovered Tg68 strain is permissive for bradyzoite differentiation and forms the foundation for the screening protocol in this study. We have developed protocols for HTS of compound libraries that target for tachyzoites and bradyzoites and further validated hits in assays against ex vivo produced bradyzoites. Several of the hits identified in this screen have precedent for being antimicrobial; however, they pose challenges for selectively leading to considerable host toxicity. The top hits identified here were potent, so potentially medicinal chemistry efforts could address the selectivity issue in future studies. Additionally, the methodology develop here offers promise for future HTS to identify potent and selective leads.

## Experimental Section

### Compounds and liquid handling

The LOPAC compound library was plated in a 384-well plate format for primary screening and determination of half-maximal effective concentration (EC_50_) of primary hits by Calibr at Scripps Research (La Jolla, CA). The plates were stored at -80°C prior to use. All liquid handling steps (host and parasite cell seeding, media exchange, compound transfer to assay plates, and addition of luciferase reagents) were carried out in a semi-automated facility to ensure efficient and consistent execution of assays across all replicates (High-Throughput Screening Center, Washington University School of Medicine). Nine top hit compounds and two control compounds, atovaquone (Sigma#A7986) and pyrimethamine (Sigma#46706), were obtained from commercial sources, and follow-up assays were manually performed in-house.

### Construction of transgenic parasites and parasite culture

Tg68 pTub1:Fluc,DHFR parasite lines were generated by utilizing a pre-existing pTub:Fluc plasmid ^33^, which provides constitutive firefly luciferase (Fluc) expression from the alpha tubulin promoter, and a pyrimethamine-resistant DHFR selectable marker. Tg68 tachyzoites were electroporated with 50 μg of the plasmid and subsequently selected using pyrimethamine (3 μM).

The plasmid pSAG1:EGFP-DHFR-BAG1:nLuc was constructed by assembling fragments encoding EGFP driven by the *SAG1* promoter, nLuc driven by the *BAG1* promoter, DHFR pyrimethamine resistance marker, and pNJ-26 vector using NEBuilder HiFi DNA Assembly Master Mix (NEB). Tg68 pBAG1:nLuc, DHFR tachyzoites (Tg68pSAG1:EGFP-DHFR-BAG1:nLuc) were electroporated with 50 μg of this plasmid and selected with pyrimethamine (3 μM) in order to establish stable parasite lines. Detailed primer information can be found in **Table S4**.

Stable clones were isolated through limiting dilution, and the expression of the transgene was confirmed by luciferase expression. Clonal transgenic Tg68 tachyzoites were maintained by serial passage in T25 flasks with confluent HFFs in D10 medium (Dulbecco’s Modified Eagle’s Medium, DMEM; Thermo Fisher, 10% fetal bovine serum (FBS), 2 mM glutamine (Sigma), and 10 μg/mL gentamicin (Thermo Fisher)) at 37°C and 5% CO_2_.

### In Vitro Assays for bradyzoite and tachyzoite growth inhibition

#### Host cell culture and seeding

Human Foreskin Fibroblasts (HFF-1 SCRC-1041, ATCC) cells were cultured in T175 flasks using D10 medium. Prior to seeding the cells to the assay plates, a cell suspension was prepared by treating with trypsin (0.5 g/l porcine trypsin and 0.2 g/L EDTA in Hank’s Balanced Salt Solution with phenol red) for 5 min. EC_50_ assays (half maximal effective concentration) were conducted in a 384-well plate format. Confluent HFF cells were seeded 4 days in advance using a Multidrop Combi dispensing 80 uL/well, while stirring the cell suspension at 350 rpm. The plates were stacked in a Cytomat rack on a level surface at room temperature until all plates were completed, and then they were placed in a 37°C incubator with 5% CO_2_. To minimize edge effects, only the inner 240 wells of each plate were utilized.

#### Bradyzoite Primary Screening and EC_50_ Assay

The medium from the HFF cell plates was aspirated using a Biotek ELx405CW, leaving 20 µL/well, before infection with parasites. Freshly harvested Tg68pBAG1:nLuc,DHFR parasites (3×10^3^) in a 40 μL volume were then added to each well (total volume of 60 μL/well, containing 0.1% DMSO in D10) using a Fluotics BXi-50F while stirring the parasite suspension at 350 rpm. The plates were incubated for 2 hr at 37°C in a 5% CO_2_ incubator to allow parasite invasion. Afterwards, the culture medium was removed and switched to either 80 μL/well of Alkaline (RPMI 1640 (Sigma, #R6504) containing 1% FBS and 50 mM HEPES (Sigma), adjusted to pH 8.2), or Glutamine (glucose-free RPMI 1640 (Sigma, #R1383) containing 1% FBS and 50 mM HEPES (Sigma), 10 mM glutamine, adjusted to pH 7.2) medium. The culture was then incubated at 37°C in ambient CO_2_ and maintained for 10 days, with media changes occurring on day 3 and 6 using a Biotek ELx405CW and Multidrop Combi inside a biohood to minimize contamination. On day 6, before the compound was added, the plates were aspirated, and 40 µL/well of fresh media was added, followed by the transfer of 40 µL/well of 2× compound using a Fluotics BXi-50F. On day 10, the plates were equilibrated to room temperature for 30 min prior to the assay readout. To read the plates, the medium was aspirated, 20 uL of Promega Nano-Glo reagent (Promega N1150) was dispensed into each well using a Multidrop Combi, and the plates were covered with a black lid. The plates were then incubated for 10 min at room temperature, and the readout was performed using Envision.

#### Tachyzoite Primary Screening and EC_50_ Assay

Before being infected with parasites, the medium was removed from the HFF cell plates using a Biotek ELx405CW, leaving 20 µL/well. Freshly harvested Tg68pTub1:Fluc parasites (2×10^4^) in a 20 μL volume were then added to each well while stirring the parasite suspension at 350 rpm, and 40 µL of 2× compound solutions were transferred (80 μL/well total volume, containing 0.1% DMSO in D10) using a Fluotics BXi-50F. The plates were incubated for 72 hr at 37 °C in a 5% CO_2_ incubator. Before the assay readout, the plates were equilibrated to room temperature for 30 min. To read the plates, the medium was aspirated to 20 µL using a Biotek ELx405CW, 20 µL of Promega Bright-Glo reagent (Promega E2650) was dispensed into each well, and the plates were covered with a black lid. The plates were then incubated for 10 min at room temperature using a Multidrop Combi, and the readout was performed using Envision.

### In Vitro Assays for Host cell cytotoxicity

Cytotoxicity against host cells host cells was tested using human hepatocellular carcinoma cells (HepG2, ATCC-HB-8065) and human monocytic tumor line (THP-1, ATCC-TIB-202), which were maintained according to ATCC recommendations. Host cell lines were tested negative for mycoplasma using an e-Myco plus kit (Intron Biotechnology). Compounds were diluted to 2× concentration, and a 10-dose serial dilution series was prepared through stepwise, 3-fold dilutions in recommended media with 0.1% DMSO. Host cells were seeded at a density of 10^4^ cells/well (100 μL vol) to achieve sub-confluent monolayers for a 72 hr growth assay. Prior to compound addition, THP-1 cells were treated with 10 ng/mL phorbol 12-myristate 13 acetate for 24 hr to facilitate differentiation into macrophages. HepG2 cells were treated with compounds 6 hr post-seeding (200 μL final volume, 0.05% DMSO) and incubated under 37 °C, 5% CO_2_ culture conditions. At 72 hr post compound addition, culture media were aspirated to 80 μL, and an equal volume of CellTiter-Glo Luminescent Cell Viability Assay reagent (Promega G7571) was added. Luciferase activity was measured using a BioTek Cytation 3 equipped with Gen5 software (v3.08). Each assay was repeated with two technical replicates within two independent biological replicates. Statistical analyses were performed using Prism 10 (GraphPad Software, Inc.). Dose-response inhibition curves for host cell toxicity (CC_50_ values) were generated using the “Log(inhibitor) vs normalized response variable slope” function. The reported values represent averages from three biological replicates.

### Ex-vivo Bradyzoite differentiation assay

CBA/CaJ mice (Strain # 000654) from Jackson Laboratory were housed in an approved facility at Washington University School of Medicine, and all animal studies followed ethical guidelines approved by the Institutional Animal Care and Use Committee. CBA/CaJ mice were infected by oral gavage with 5-10 tissue cysts from the brain homogenate of previously infected mice. The brains of CBA/CaJ mice infected with the TgME49-EW strain ^41^ were collected at 1-2 mos post-infection, homogenized, and tissue cysts were isolated using Percoll gradients, as described previously ^54^. To release the bradyzoites from the tissue cysts, purified tissue cysts were treated with an acid-pepsin solution (170 mM NaCl, 60 mM HCl), and a freshly prepared pepsin solution (0.1 mg/mL in 1xPBS) for 10 min at 37 °C. The reaction was stopped by adding a neutralization buffer (94 mM Na_2_CO_3_). The liberated bradyzoites were then evenly distributed into a duplicate set of 6-well plates (technical replicates), each containing 5 mL of culture media. Each plate included a negative control (media with 0.1% DMSO), the previously characterized PheRS inhibitor BRD7929 (0.5 µM)^33^, and pyrimethamine (2.0 µM). Compounds were tested at EC_90_ and 3XEC_90_ concentrations based on their corresponding in vitro tachyzoite growth inhibition (TgEC_50_) assays (Table S3). After a 4 hr treatment, the compounds were removed from one set of plates by washing with 3 times in PBS and replacement with compound-free D10 medium. The plates were then incubated at 37°C, 5% CO_2_ undisturbed for 12-14 days to form plaques. Plaque quantification was performed by fixing the plates in 100% ethanol for 5 min at room temperature, followed by staining with a 0.1% crystal violet solution for 10 min. After rinsing with water and air drying, plaque quantification was conducted using a Nikon eclipse ts2 microscope equipped with a 4X objective. The number of plaques from two biological replicates was normalized to DMSO control as a percentage.

### Statistical analysis

Statistical comparisons were performed in Prism (GraphPad). Data were first analyzed for normal distribution and according to the outcome, were then non-parametric tests applied. Statistical test and *P* values are given in the figure legends.

## Acknowledgments

We thank Jennifer Barks for HFF cell culture and for providing administrative support for research materials. We acknowledge Maxene Ilagan and Mike Prinsen from Washington University’s High Throughput Screening Center for their assistance with assay set-up and data handling. Furthermore, we thank Kaycie Morwood and her team from Calibr – A Division of Scripps Research for their assistance with compound management and plating. The ME49EW was a kindly provided by Michael White, University of South Florida. Partial support provided by an NIH grant (AI143857) and a pilot grant from the Center for Drug Discovery at Washington University (PJ00027513).

## Abbreviations used

DMSO: Dimethylsulfoxide
DHFR: Dihydrofolate Reductase
EDTA: Ethylenediaminetetraacetic acid
HFF: Human Foreskin Fibroblasts
HepG2: human hepatocellular carcinoma cell
THP-1: human monocytic tumor cell
HTS: High Throughput Screening
nLuc: Nano Luciferase
Fluc: Firefly Luciferase
PheRS: Phenylalanine tRNA synthetase

## Supplemental Materials

**Table S1 HTS parameters for lines expressing luciferase**.

**Table S2 HTS parameters for the LOPAC screen**.

**Table S3 Activity of Primary Hits in growth inhibition assays**.

**Table S4 Primers used in the study**.

---

**Figure S1.**
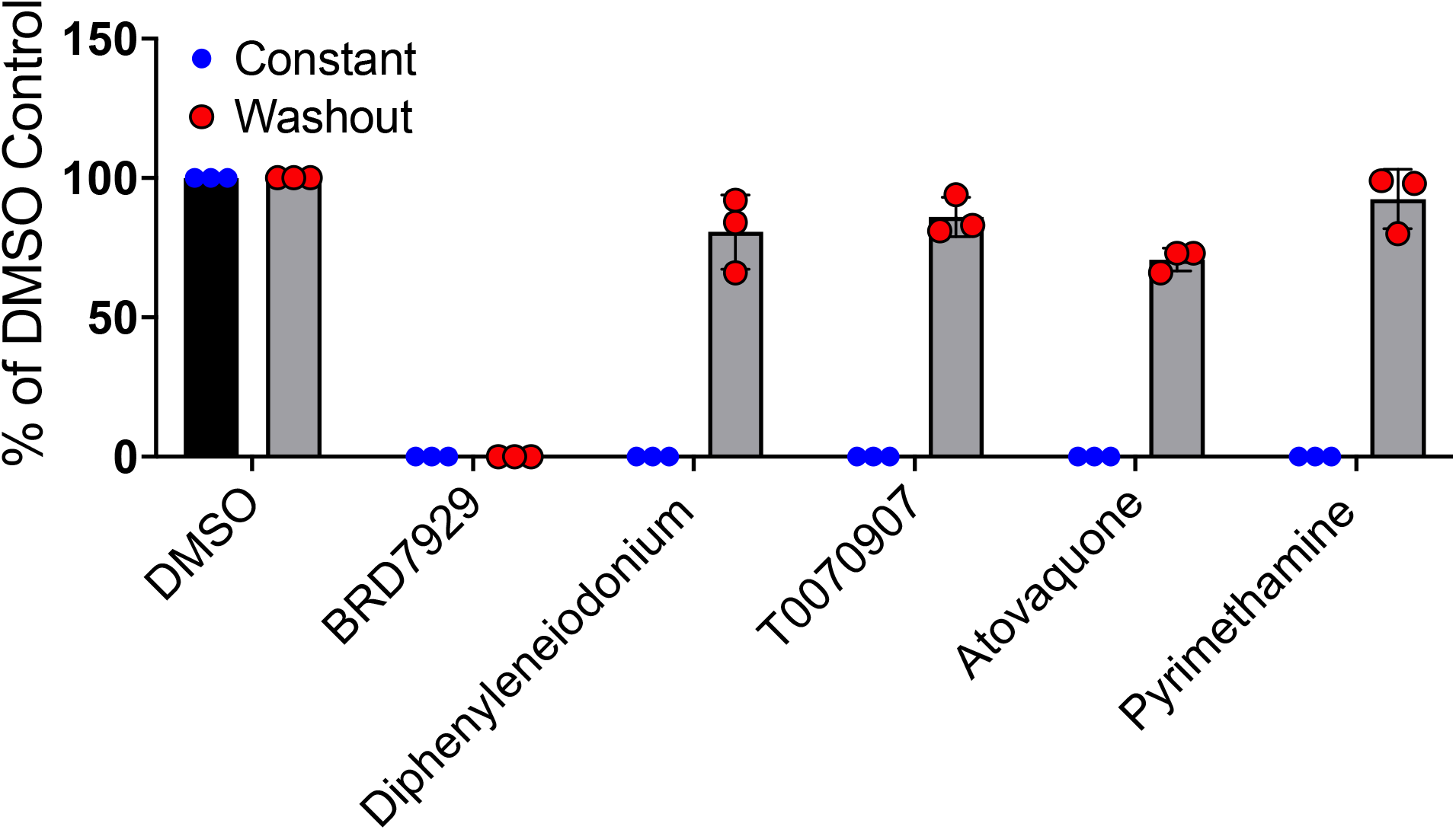
Testing of additional TOP HIT compounds for activity against ex vivo bradyzoites. Plaque number from 3 biological replicates for Diphenyleneiodonium, T0070907 and reference compounds in continuous treatment vs. 4 hr treatment and washout presented in average percentage with standard deviation bar.

